# A sensitized mutagenesis screen in Factor V Leiden mice identifies novel thrombosis suppressor loci

**DOI:** 10.1101/080432

**Authors:** Randal J. Westrick, Kärt Tomberg, Amy E. Siebert, Guojing Zhu, Mary E. Winn, Sarah L. Dobies, Sara L. Manning, Marisa A. Brake, Audrey C. Cleuren, Linzi M. Hobbs, Lena M. Mishack, Alexander Johnston, Emilee Kotnik, David R. Siemieniak, Jishu Xu, Jun Z. Li, Thomas L. Saunders, David Ginsburg

**Affiliations:** Oakland University Department of Biological Sciences; Oakland University Center for Data Science and Big Data Analysis; Life Sciences Institute, University of Michigan; Bioinformatics and Biostatistics Core, Van Andel Research Institute; Department of Human Genetics, University of Michigan; Howard Hughes Medical Institute, University of Michigan; Transgenic Animal Model Core, University of Michigan; Departments of Internal Medicine and Pediatrics, University of Michigan

**Keywords:** Venous thromboembolism, Factor V Leiden, ENU mutagenesis

## Abstract

Factor V Leiden (*F5^L^*) is a common genetic risk factor for venous thromboembolism in humans. We conducted a sensitized ENU mutagenesis screen for dominant thrombosuppressor genes based on perinatal lethal thrombosis in mice homozygous for *F5^L^* (*F5^L/L^*) and haploinsufficient for tissue factor pathway inhibitor (*Tfpi*^+/−^). *F8* deficiency enhanced survival of *F5^L/L^ Tfpi*^+/−^ mice, demonstrating that *F5^L/L^ Tfpi*^+/−^ lethality is genetically suppressible. ENU-mutagenized *F5^L/L^* males and *F5^L/+^ Tfpi*^+/−^ females were crossed to generate 6,729 progeny, with 98 *F5^L/L^ Tfpi*^+/−^ offspring surviving until weaning. Sixteen lines exhibited transmission of a putative thrombosuppressor to subsequent generations, with these lines referred to as *MF5L* (<u>**M**</u>odifier of <u>**F**</u>actor <u>**5 L**</u>eiden) 1-16. Linkage analysis in *MF5L6* identified a chromosome 3 locus containing the tissue factor gene (*F3*). Though no ENU-induced *F3* mutation was identified, haploinsufficiency for *F3* (*F3*^+/−^) suppressed *F5^L/L^ Tfpi*^+/−^ lethality. Whole exome sequencing in *MF5L12* identified an *Actr2* gene point mutation (p.R258G) as the sole candidate. Inheritance of this variant is associated with suppression of *F5^L/L^ Tfpi*^+/−^ lethality (p=1.7x10^−6^), suggesting that *Actr2*^p.R258G^ is thrombosuppressive. CRISPR/Cas9 experiments to generate an independent *Actr2* knockin/knockout demonstrated that *Actr2* haploinsufficiency is lethal, supporting a hypomorphic or gain of function mechanism of action for *Actr2*^p.R258G^. Our findings identify *F8* and the *Tfpi/F3* axis as key regulators in determining thrombosis balance in the setting of *F5^L^* and also suggest a novel role for *Actr2* in this process.

**Significance Statement:** Venous thromboembolism (VTE) is a common disease characterized by the formation of inappropriate blood clots. Inheritance of specific genetic variants, such as the Factor V Leiden polymorphism, increases VTE susceptibility. However, only ~10% of people inheriting Factor V Leiden develop VTE, suggesting the involvement of other genes that are currently unknown. By inducing random genetic mutations into mice with a genetic predisposition to VTE, we identified two genomic regions that reduce VTE susceptibility. The first includes the gene for blood coagulation Factor 3 and its role was confirmed by analyzing mice with an independent mutation in this gene. The second contains a mutation in the Actr2 gene. These findings identify critical genes for the regulation of blood clotting risk.

## Introduction

Venous thromboembolism (VTE) is a common disease that affects 1 to 3 per 1000 individuals per year(1). VTE susceptibility exhibits a complex etiology involving contributions of both genes and environment. Genetic risk factors explain approximately 60% of the overall risk for VTE(2). Recent large-scale genome-wide association studies (GWAS) confirmed *ABO*, *F2 F5*, *F11*, *FGG* and *PROCR* as thrombosis susceptibility genes, with only two additional novel loci, TSPAN15 and SLC44A2 identified(3–6), leaving the major component of VTE genetic risk still unexplained.

The Factor V Leiden variant (*F5^L^*) is a common inherited risk factor for VTE with an average allele frequency of 3.5% in the European population(7–9). *F5^L^* is estimated to account for up to 25% of the genetically-attributable thrombosis risk in these populations(7). However, penetrance is incomplete, with only ~10% of *F5^L^* heterozygotes developing thrombosis in their lifetimes. The severity of thrombosis also varies widely among affected individuals(10), limiting the clinical utility of *F5^L^* genotyping in the management of VTE(11).

The incomplete penetrance and variable expressivity of thrombosis among *F5^L^* patients can at least partially be explained by genetic interactions between *F5^L^* and other known thrombotic risk factors such as hemizygosity for antithrombin III or proteins C or S, as well as the common prothrombin 20210 polymorphism(10, 12, 13). However, <2% of *F5^L^* heterozygotes would be expected to co-inherit a mutation at one or more of these loci, suggesting that a large number of additional genetic risk factors for VTE and/or modifiers of *F5^L^* remain to be identified(3, 10).

Mice carrying the orthologous *F5^L^* mutation exhibit a mild to moderate prothrombotic phenotype closely mimicking the human disorder(14). We previously reported a synthetic lethal interaction between *F5^L^* homozygosity (*F5^L/L^*) and hemizygosity for tissue factor pathway inhibitor (*Tfpi*^+/−^)(15). Nearly all mice with this lethal genotype combination (*F5^L/L^ Tfpi*^+/−^) succumb to widespread, systemic thrombosis in the immediate perinatal period(15).

ENU mutagenesis in mice has been used effectively to identify novel genes involved in a number of biological processes(16, 17). ENU-induced germline mutations transmitted from a mutagenized male mouse (G0) occur at ~1.5 mutations per megabase, at least 50 fold higher than the endogenous background mutation rate(18). Several previous reports have successfully applied an existing phenotype as a sensitizer to identify modifier genes. A dominant suppressor screen in *Mecp2* deficient mice (Rett syndrome) identified a mutation in squalene epoxidase (*Sqle)* as a heritable suppressor, resulting in prolonged survival and amelioration of neurologic manifestations(19). Other successful sensitized screens include analysis of mouse mutants predisposed to diabetic nephropathy(20), a screen in *Sox10* haploinsufficent mice identifying the *Gli3* gene as a modifier of neurochristopathy(21) and identification of a mutation in the *c-Myb* gene as a dominant modifier for platelet count in *Mpl* deficient mice (congenital thrombocytopenia)(22). We now report the results of a dominant, sensitized ENU mutagenesis screen for suppressors of *F5^L/L^ Tfpi*^+/−^ dependent lethal thrombosis.

## Results and Discussion

### *F8* deficiency suppresses *F5^L/L^ Tfpi*^+/−^ lethality

X-linked hemophilia A results in a moderate to severe bleeding disorder in humans and is caused by mutations in the *F8* gene. To test whether the *F5^L/L^ Tfpi*^+/−^ lethal thrombotic phenotype is suppressible by hemophilia A in mice, triple heterozygous *F5^L/+^ Tfpi^+/−^ F8*^+/−^ female mice were generated and crossed to *F5^L/L^* male mice (Fig. 1A). One quarter of conceptuses are expected to carry the *F5^L/L^ Tfpi*^+/−^ genotype, with half of the total expected male conceptuses completely *F8* deficient (*F8^−^*). Thus, 1/16^th^ of the overall offspring from this mating are expected to be *F5^L/L^ Tfpi^+/−^ F8^−^* (males). Similarly, 1/16^th^ of the progeny should be *F5^L/L^ Tfpi^+/−^ F8*^+/−^ (females). A total of 163 progeny from this cross were genotyped at weaning, resulting in 8 *F5^L/L^ Tfpi^+/−^ F8^−^* male mice observed (and 0 *F5^L/L^ Tfpi^+/−^ F8^+^*, p=0.02) and 2 *F5^L/L^ Tfpi^+/−^ F8*^+/−^ female mice (and 1 *F5^L/L^ Tfpi^+/−^ F8^+/+^*, p=0.9). These results demonstrate that *F5^L/L^ Tfpi*^+/−^ thrombosis is genetically suppressible by *F8* deficiency with nearly complete penetrance in *F8^−^* male mice and are consistent with human studies demonstrating *F8* level as an important VTE risk factor(23).

**Figure 1:**
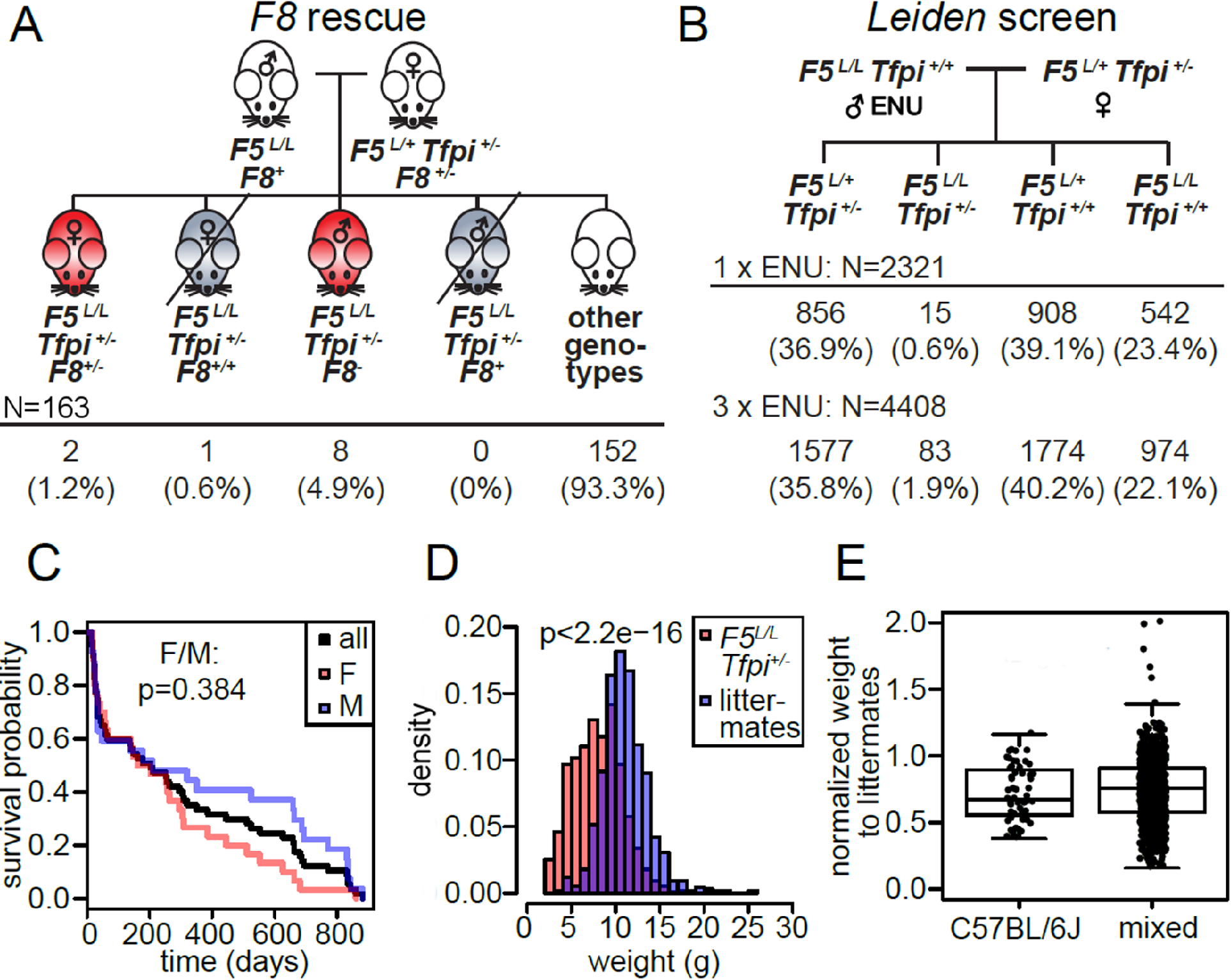
*F8* deficient thrombosuppression and design of the Leiden ENU mutagenesis screen. **A.** The mating scheme and observed distributions of the *F5^L/+^ Tfpi^+/−^ F8* deficiency rescue experiments. *F8^−^* results in suppression of the *F5^L/L^ Tfpi*^+/−^ phenotype. **B.** The mating scheme and observed distribution of the Leiden screen. *F5^L/L^ Tfpi^+/+^* male mice were mutagenized with either 1 x 150mg/kg or 3 x 90 mg/kg ENU and bred with non-mutagenized *F5^L/+^ Tfpi*^+/−^ females. Fifteen and 83 *F5^L/L^ Tfpi*^+/−^ progeny, respectively were observed in each of the dosing regimens, with over twice the rate of *F5^L/L^ Tfpi*^+/−^ survivors in the progeny of the 3 x 90 mg/kg treated mice. **C.** There was no significant difference in survival between male and female *F5^L/L^ Tfpi*^+/−^ putative suppressor mice, (p=0.384). Normal weaning and breeding ages are 20 days and 42 days, respectively. **D.** *F5^L/L^ Tfpi*^+/−^ putative suppressor mice were significantly smaller than their non-*F5^L/L^ Tfpi*^+/−^ littermates. **E.** *F5^L/L^ Tfpi*^+/−^ putative suppressors were smaller than their littermates of other genotypes (p<8.8 x 10^−12^ for B6, and p=2.2 x 10^−16^ for mixed B6-129S1) regardless of whether they were on the pure B6 or mixed B6-129S1 genetic backgrounds (p=0.327 between B6 and mixed backgrounds).

### The *F5^L/L^ Tfpi*^+/−^ phenotype is suppressed by dominant ENU induced mutations

A sensitized, genome-wide ENU mutagenesis screen for dominant thrombosis suppressor genes was implemented as depicted in Figure 1B. ENU mutagenized G0 *F5^L/L^* males were crossed to *F5^L/+^ Tfpi*^+/−^ females to generate G1 mice, which were screened by genotyping at weaning for *F5^L^* and *Tfpi*^+/−^. Previously described visible dominant mutant phenotypes(24), including belly spotting and skeletal abnormalities, were observed in approximately 5.9% of G1 offspring, similar to the ~4.2% rate of observable mutants in previous studies(24). This is consistent with the ~20-30 functionally significant mutations per G1 mouse expected with this ENU mutagenesis protocol(25). Although 25% of G1 embryos from this cross are expected to carry the synthetic lethal *F5^L/L^ Tfpi*^+/−^ genotype, most are lost at birth. Given a total of 6631 G1s for the other 3 genotypes observed at weaning (~1/3 for each), a similar number of *F5^L/L^ Tfpi*^+/−^ G1 conceptuses, ~2210 (6631 X 1/3), would have been expected. The 98 live *F5^L/L^ Tfpi*^+/−^ mice (45 females, 53 males) thus represented 4.4% of the expected with this genotype. Survival data were collected for 57 of the *F5^L/L^ Tfpi*^+/−^ G1 mice, with 34 living past 70 days of age (precise dates of death were not available for the remaining 41 mice). No significant sex-specific differences in survival were observed (Fig. 1C).

Heritability for each of the 44 G1 putative suppressor mutants who lived to breeding age was evaluated by a progeny test backcross to C57BL/6J (B6) *F5^L/L^* mice. The observation of one or more *F5^L/L^ Tfpi*^+/−^ offspring surviving to weaning increased the likelihood that a particular Modifier of Factor 5 Leiden (*MF5L*) line carries a transmissible suppressor mutation. Out of the original 98 surviving *F5^L/L^ Tfpi*^+/−^ G1 mice, 75 produced no offspring surviving to weaning, either due to infertility or the above mentioned early lethality, with >50% of these mice (37 of 75) exhibiting a grossly runted appearance. Approximately half of the *F5^L/L^ Tfpi*^+/−^ G1 mice that attained breeding age (23/44) produced 1 or more G2 progeny surviving to weaning, 7 (2 males and 5 females) produced no *F5^L/L^ Tfpi*^+/−^ G2s, including 4 G1s with 8 or more offspring of other genotypes. Sixteen *F5^L/L^ Tfpi*^+/−^ G1 mice produced one or more *F5^L/L^ Tfpi*^+/−^ progeny when bred to B6 *F5^L/L^* mice (see Methods). These 16 potential thrombosuppressor mouse lines are designated *MF5L1-16*. The number of total progeny, genotypic distribution, and penetrance of the *F5^L/L^ Tfpi*^+/−^ mice in each line are listed in Table 1. Within these suppressor lines, mice with the *F5^L/L^ Tfpi*^+/−^ genotype were ~30% smaller than their *F5^L/L^* littermates at the time of weaning (p<2.2x10^−16^, Fig. 1D), with this difference maintained after outcrossing to the 129S1 strain (Fig. 1E).

**Table 1:**
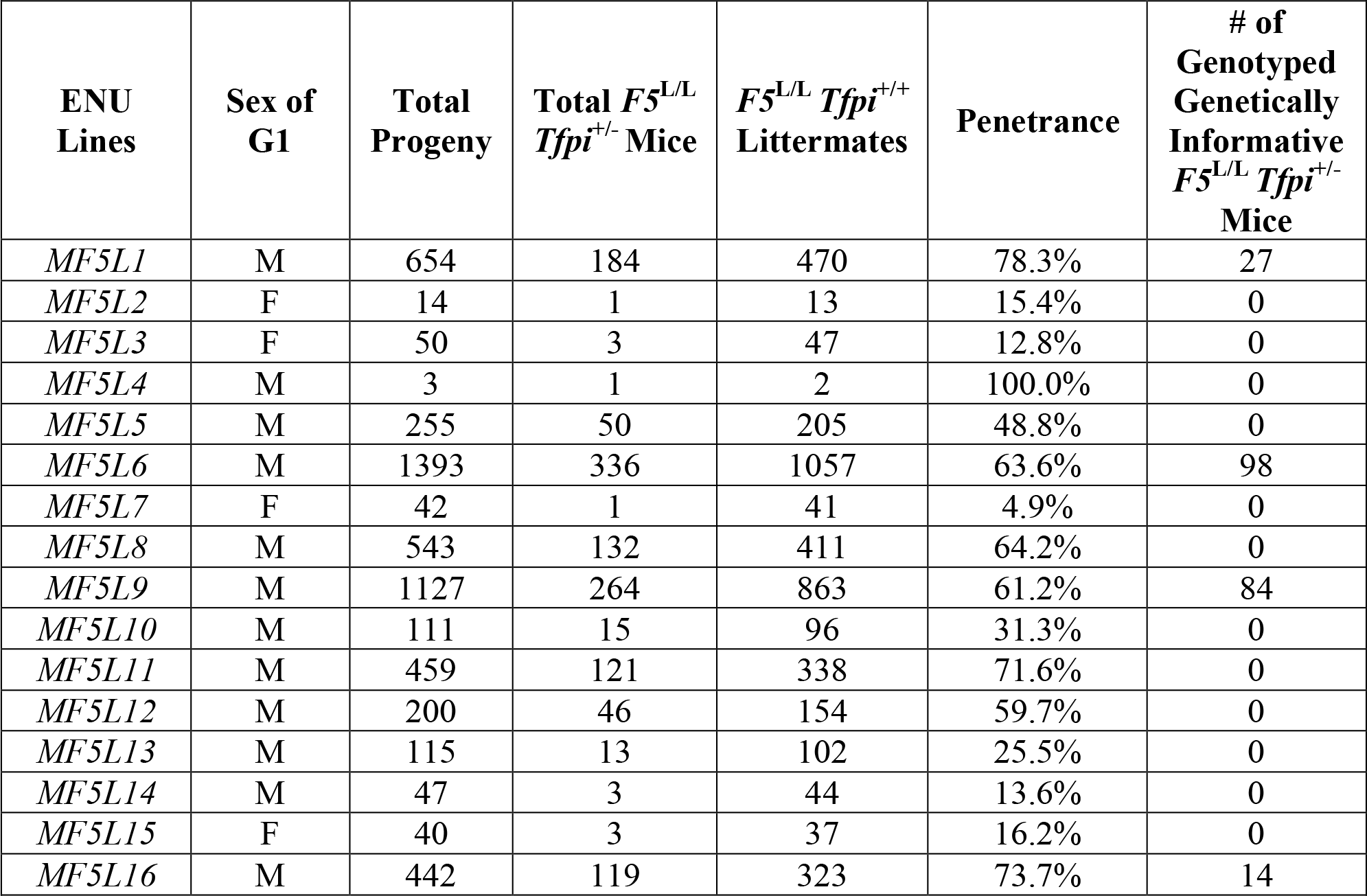
Progeny Genotypes and Penetrance of Putative *MF5L* Thrombosuppressor Genes

Penetrance was calculated as total *F5*^L/L^ *Tfpi*^+/−^ divided by ½ of the number of *F5*^L/L^ *Tfpi*^+/+^ littermates.

Previous reports based on gene function in the specific locus test estimate an ENU-induced mutation rate of 1/700 loss of function mutations per locus for the ENU dosing regimen used here(26). This mutation rate predicts that our screen of 6,729 G1 progeny (2,210 *F5^L/L^ Tfpi*^+/−^ expected) should have produced ~3 mutations per gene averaged over the entire genome, with 54% of these mutations expected to be null(16), resulting in 1.5X genome coverage for loss of function mutations.

### The *MF5L6* suppressor mutation maps to a chromosome 3 interval containing *F3*

In order to map putative ENU-induced suppressor mutations, surviving *F5^L/L^ Tfpi*^+/−^ mice were intercrossed with *F5^L/L^* mice that had been backcrossed onto the 129S1/SvIMJ strain (129S1). Crosses between *F5^L/L^* and *F5^L/+^ Tfpi*^+/−^ mice (both *F5^L^* and *Tfpi^−^* backcrossed > 12 generations onto 129S1) confirmed the lethality of the *F5^L/L^ Tfpi*^+/−^ genotype on the 129S1 background (Table S1). The 4 lines containing the largest number of genetically informative B6-129S1 mixed background *F5^L/L^ Tfpi*^+/−^ offspring (*MF5L1, 6, 9* and *16*) were used for gene mapping. Though the *MF5L1*, *MF5L9* and *MF5L16* lines were successfully expanded to pedigrees containing 27, 84, and 14 *F5^L/L^ Tfpi*^+/−^ informative offspring, respectively, genotyping for a total of ~800 markers in each cross failed to identify any loci with a LOD ≳3 (maximum LODs for *MF5L1=1.15*, *MF5L9=2.5* and *MF5L16=1.61*). Though we cannot exclude cosegregation of more than one suppressor mutation, the absence of a clear linkage signal for each of these lines likely reflects complex mouse strain modifier gene interactions, which are known to significantly impact mouse phenotypes(10, 27) and confound linkage analysis(28). Consistent with this hypothesis, we have previously documented poorer survival to weaning in mixed B6-129S1 *F5^L/L^* mice compared to littermates(14). We extended these observations by the analysis of additional *F5^L/+^* and *F5^L/L^* littermates, with mice of the *F5^L/L^* genotype demonstrating a 50% reduction in survival in the 129S1 versus B6 strain backgrounds (Table S1).

*MF5L*6 was maintained for 12 generations on both the mixed and B6 backgrounds and produced a total of 336 *F5^L/L^ Tfpi*^+/−^ mice (98 on the mixed B6-129S1 background and therefore useful for linkage analysis, See Table 1). Genome-wide SNP genotyping was performed on DNA from these 98 genetically informative *F5^L/L^ Tfpi*^+/−^ mice, with multipoint linkage analysis shown in Figure 2A. Since the genetic intervals around the *F5* and *Tfpi* loci cannot be accurately assessed for linkage, these regions of chromosomes 1 and 2 were excluded from linkage analysis (See Fig. 2A Legend and Methods). A single locus with a significant LOD score of 4.49 was identified on chromosome (Chr) 3, with the 1 LOD interval (117.3-124.8Mb) containing 43 refseq annotated genes (Fig. 2B).

**Figure 2:**
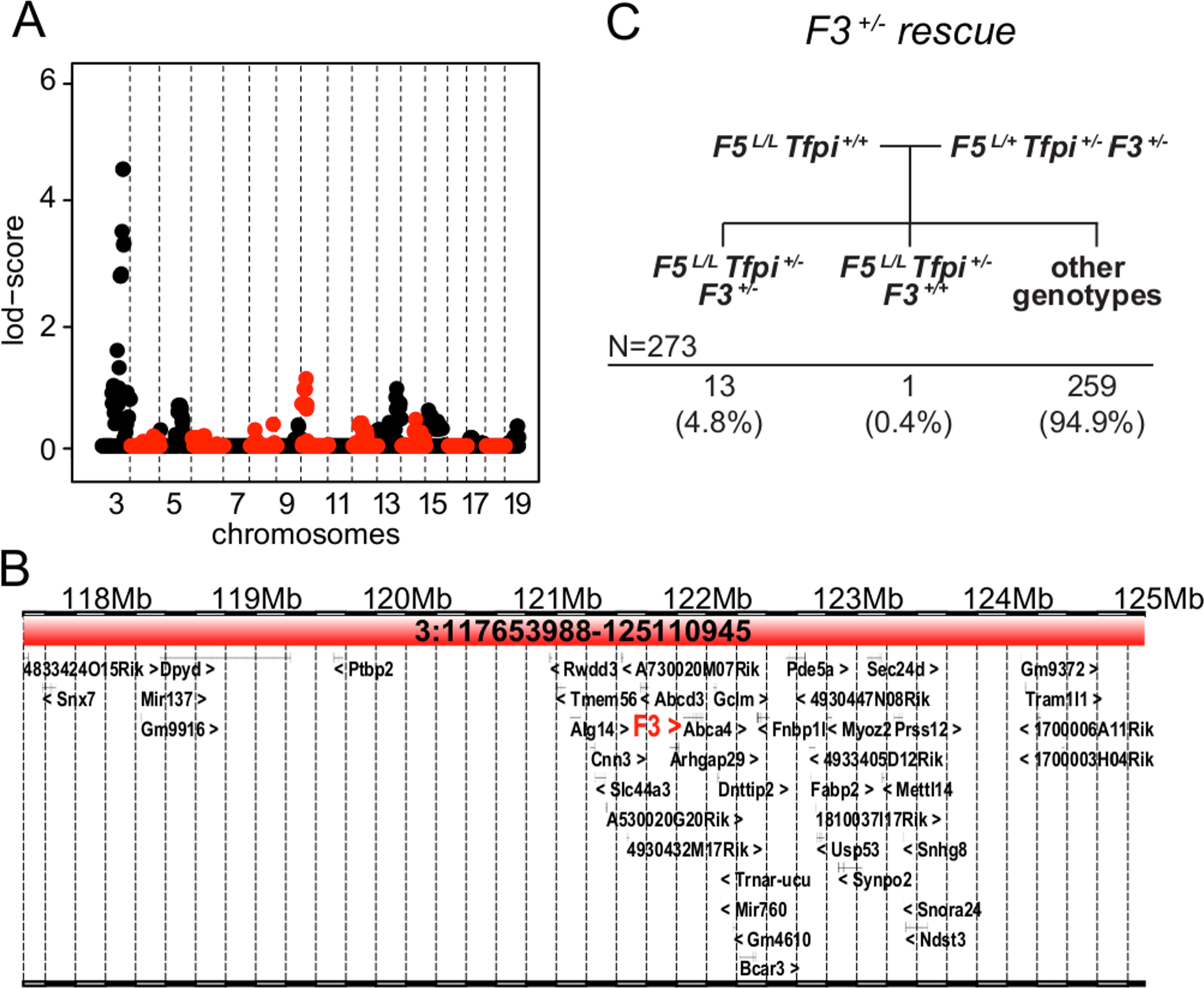
The *MF5L6* suppressor locus maps to Chr3. **A.** Linkage analysis for the *MF5L6* line. Alternating red and black is used to highlight the chromosomes. Chrs 1 and 2 were excluded from further analysis since they contain the *F5* and *Tfpi* genes, whose segregation was restricted by required genotypes at these loci. The Chr3 peak had the highest LOD score in the Chr3 subregion:117.3-124.8Mb (maximum LOD=4.49, 1 LOD interval, significance threshold of LOD >3.3(49)). **B.** The Chr3 candidate interval (Chr3:117.3-124.8 Mb) contains 43 refseq annotated genes, including *F3*. **C.** The mating scheme and observed distribution of offspring to test *F3* deficiency as a suppressor of *F5^L/L^ Tfpi*^+/−^. *F3*^+/−^ results in incompletely penetrant suppression of the *F5^L/L^ Tfpi*^+/−^ phenotype.

The *F3* gene located within this interval (Chr3:121.7 Mb) (Fig. 2B) encodes tissue factor (TF), a procoagulant component of the hemostatic pathway that has *Tfpi* as its major regulator. Quantitative or qualitative deficiencies in *F3* are thus highly plausible candidates to suppress the *F5^L/L^ Tfpi*^+/−^ phenotype. To test *F3* as a candidate suppressor of the *F5^L/L^ Tfpi*^+/−^ phenotype, an independent *F3* null allele was introduced and triple heterozygous *F5^L/+^ Tfpi^+/−^ F3*^+/−^ mice were crossed to *F5^L/L^* B6 mice (Fig. 2C). Of 273 progeny genotyped at weaning, 13 *F5^L/L^ Tfpi*^+/−^ *F3*^+/−^ were observed (and 1 *F5^L/L^ Tfpi^+/−^ F3^+/+^*, p=9.7x10^−5^). We also observed significantly fewer male than female *F5^L/L^ Tfpi^+/−^ F3*^+/−^ mice (2 vs. 11 p=0.03). Thus, haploinsufficiency for *F3*^+/−^ suppresses the synthetic lethal *F5^L/L^ Tfpi*^+/−^ phenotype with incomplete penetrance (33%) that also differs by sex (10% for males and 67% for females). In contrast, the *MF5L6* line exhibited an overall penetrance of 72.4%, with similar male/female penetrance. Gender-specific differences in venous thrombosis rates have previously been reported, including contributions from oral contraceptives and hormone replacement therapy(29–31). This difference in penetrance could be due to 129S1 strain effects in the *MF5L6* line or differences between a *F3* regulatory mutation in *MF5L6* compared to the *F3* loss of function allele used here.

Whole exome sequencing data analysis of a F5L/L Tfpi+/- mouse from MF5L6 failed to identify an ENU variant in F3 or in any other genes in the nonrecombinant interval, or more broadly on the entire Chr3. This is a particularly gene rich region (Figure 2B), and errors in annotation could obscure the responsible variant. Of note, this interval also includes *Slc44a3*, a paralog of *Slc44a2*, the latter previously identified as a potential modifier of VTE risk in humans(6). Although additional ENU variants were identified on other chromosomes, none co-segregated with the survival phenotype in line *MF5L6* (Table S2). Sanger sequencing analysis of the full set of *F3* exons and introns, as well as 5kb upstream of exon 1, also failed to identify an ENU-induced mutation. In addition, analysis of *F3* mRNA levels in liver, lung and brain tissues of adult mice failed to identify any differences in the level of expression from the ENU-mutant compared to the wildtype allele (Fig. S1).

Taken together, these data suggest that an ENU-induced *F3* regulatory mutation outside of the sequenced segment may be responsible for thrombosuppression in *MF5L6*, although we cannot exclude a regulatory mutation in another gene. Nonetheless, our findings demonstrate that *F3*/*Tfpi* balance plays a key role in thrombosis in the mouse, particularly in the setting of *F5^L^*, and suggest that modest variations in either *F3* or *Tfpi* could be important modifiers of VTE susceptibility in humans.

### Whole exome sequencing identifies candidate ENU suppressor variants for 8 *MF5L* lines

Whole exome-next generation sequencing (NGS) was performed on genomic DNA from an index *F5^L/L^ Tfpi*^+/−^ mouse (from the G2-G5 generation) from each of 8 *MF5L* lines, including the 4 lines described above, as well as 4 additional lines with large pedigrees (*MF5L5*, *MF5L8*, *MF5L11*, *MF5L12*). The mean coverage of sequenced exomes was more than 90X, with >97% of the captured region covered with at least 6 independent reads (Table S3). A total of 125 heterozygous variants were identified as candidate suppressor mutations, with 79 variants affecting protein sequence (Table S2). Of the total mutations, 54.4% were nonsynonymous single nucleotide variants (SNVs), followed by UTR (17.6%), synonymous (14.4%) and stopgain SNVs (7.2%), with the remainder being comprised of indels, splicing, and stoploss mutations. The most common mutation events were A/T→G/C transitions (35.3%), while C/G→G/C transversions were the least represented (2.5%). This spectrum of mutations is consistent with previously published ENU experiments(32). Variants exhibiting no recombination with the *Tfpi* locus on Chr2 (17 variants) were excluded from further analysis (See Methods). Sanger sequencing confirmation was performed for 62 variants, including all nonsynonymous and stopgain mutations. These variants were then checked for parent of origin (either the G1 mutagenized progeny or its non-mutagenized mate) as well as the original mutagenized G0 male. Forty-seven of these variants were identified in the G1 mouse but not in the G0 or non-mutagenized parent, consistent with ENU-induced mutations. The remaining 15 mutations were either NGS sequencing errors (11/15), de novo (2/15), or transmitted from the non-mutagenized parent (2/15) (Table S2).

Each SNV was analyzed in additional *MF5L* mice from the line in which it was identified. None of the thrombosuppressive exonic ENU-induced variants identified in lines *MF5L1, 5, 6, 8, 9, 11* and *16* segregated with the lethal phenotype as tested by Kaplan-Meier analysis using a significance threshold of p<0.05(33). Of the 7 candidate ENU-induced SNVs identified from whole exome sequence analysis for the *MF5L12* line, 1 was an NGS sequencing error and 6 were validated by Sanger sequencing as consistent with ENU-induced mutations in the G0 mice (Table S2). For each of these 6 SNVs, co-segregation with the survival phenotype was tested by Kaplan-Meier analysis of the first 31 *F5^L/L^ Tfpi*^+/−^ mice generated from the *MF5L12* line. Only one variant, a nonsynonymous SNV in the *Actr2* gene (c.772C>G, p.R258G, *Actr2^G^*), demonstrated a significant survival advantage when co-inherited with the *F5^L/L^ Tfpi*^+/−^ genotype (p=1.7×10^−6^) (Fig. 3A).

**Figure 3:**
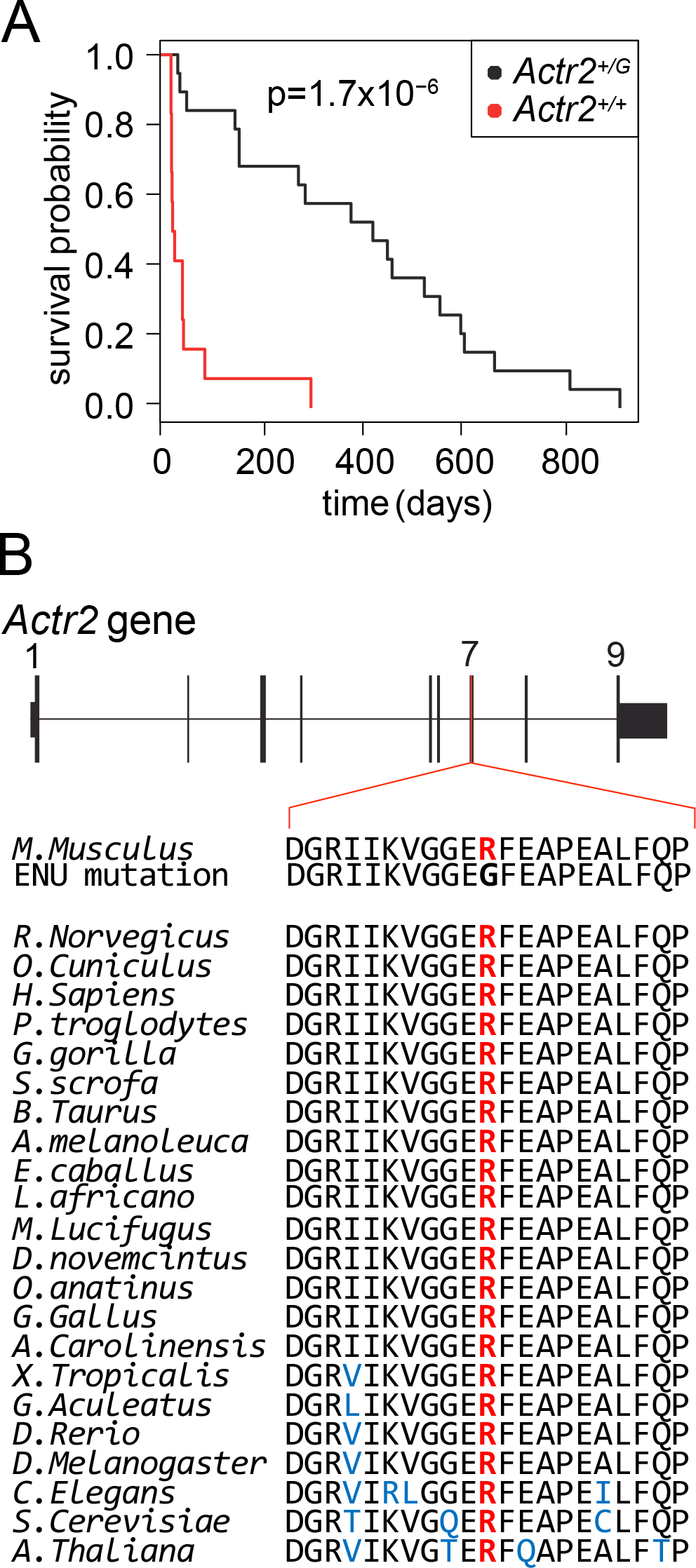
The *Actr2* R258G ENU-induced mutation is a potential thrombosis suppressor gene. **A.** Kaplan-Meier survival plot for *F5^L/L^ Tfpi*^+/−^ mice with and without the *Actr2^G^* mutation. *F5^L/L^ Tfpi^+/−^ Actr2^+/G^* mice exhibit significantly better survival than *F5^L/L^ Tfpi^+/−^ Actr2^+/+^* littermates (n=19 for *Actr2^+/G^*, n=12 for *Actr2^+/+^*, n=31 total). **B.** ARP2 amino acid R258 is highly conserved in animals, plants and fungi.

### *Actr2* as a thrombosuppressor gene

The gene *Actr2*, ARP2 actin related protein 2, encodes the ARP2 protein, which is an essential component of the Arp2/3 complex(34). ARP2, along with ARP3 and five other independent protein subunits (ARPC1-5), forms the evolutionarily conserved seven-subunit Arp2/3 complex(35). Arp2/3 is a major component of the actin cytoskeleton and is found in most eukaryotic cells including platelets(36). Arp2/3 binds to the sides of actin filaments and initiates the growth of a new filament, leading to the creation of branched actin networks that are important for many cellular processes(37). Loss of Arp2/3 function can have severe consequences, as illustrated by the embryonic lethality of mice homozygous for an ARP3 hypomorph(38). In hemostasis, the Arp2/3 complex is necessary for actin-dependent platelet cytoskeletal remodelling events, which are essential for platelet activation and degranulation(39–41). The *Actr2^+/G^* mutation results in a p.R258G substitution in exon 7 of *Actr2*, at a highly conserved amino acid position, with arginine present at this position for all 60 available vertebrate sequences (https://genome.ucsc.edu), as well as in plants and fungi (Fig. 3B). In addition, no variants at this position have been identified to date in over 120,000 human alleles(42).

### *Actr2* hemizygosity is incompatible with survival

We attempted to generate an independent *Actr2* knockin (*Actr2^G^*) allele by CRISPR/Cas9 genome editing (Fig. S2, Fig. S3, and Methods). Though highly efficient gene targeting was observed in blastocysts (Fig. S3), transfer of 275 injected embryos into foster mothers resulted in no surviving pups with a successfully targeted *Actr2* allele. These data suggest that heterozygous loss of function for *Actr2* may be incompatible with survival to term. Consistent with this hypothesis, human sequencing data from the Exome Aggregation Consortium (ExAC), which includes 60,706 individual exomes, reports a loss of function intolerance for *ACTR2* of 0.997(42). *ACTR2* mutations have not been previously associated with human disease (https://omim.org/entry/604221)(43), again consistent with early embryonic lethality. In addition, out of 373,692 mouse ENU-induced mutations listed in the Mutagenetix website, only 16 are located in the *Actr2* gene, with no predicted loss of function mutations (https://mutagenetix.utsouthwestern.edu/)(44). Taken together, these data strongly suggest that haploinsufficiency for *Actr2* is not tolerated in humans or mice. The viability of *Actr2^+/G^* mice suggests that the *Actr2^G^* allele is either hypomorphic or a unique gain of function mutation distinct from simple haploinsufficiency. Similarly, analysis of ExAC data suggests that 4 of the 6 other members of the Arp2/3 complex are intolerant to heterozygous loss of function in humans(42). Our findings suggest that subtle alterations in *Actr2* function, and potentially in other components of the actin cytoskeleton, could alter hemostatic balance and play a previously unappreciated role in thrombosis susceptibility.

The identification of novel factors involved in the regulation of hemostasis is challenging, as genes leading to marked shifts in hemostatic balance resulting in either severe bleeding or thrombosis are straightforward to identify clinically in humans, whereas subtle shifts are likely to escape detection given the multiple layers of buffering built into the complex hemostatic system(10). Homozygous deficiency (which would not be tested by our dominant suppressor strategy) for a number of hemostatic factors results in clinical bleeding, whereas heterozygous carriers remain asymptomatic. Though a single mutation in the X-chromosomal Factor VIII (or IX) gene produces severe bleeding in humans and rescues the lethal *F5^L/L^ Tfpi*^+/−^ mouse phenotype (Figure 1A), a F8 gene mutation would not be transmitted from the ENU-mutagenized male-to-male offspring and thus would be undetected. Indeed, the dominant sensitized suppressor screen reported here was undertaken to identify genes for which modest (<50%) reduction in function would significantly shift overall hemostatic balance. Such loci represent likely candidates for common human variation contributing to thrombosis and bleeding disorders. Gene variants with subtle yet significant antithrombotic effects represent attractive therapeutic targets because of a potentially wide therapeutic window with few unintended side effects. The finding of 98 *F5^L/L^ Tfpi*^+/−^ mice carrying putative thrombosis suppressor mutations (at an estimated 1.5X genome coverage) suggests that subtle alterations at a number of loci are capable of suppressing the *F5^L/L^ Tfpi*^+/−^ lethal thrombotic phenotype. The complex strain-specific genetic modifiers that confounded the genetic linkage analysis are consistent with this model. Nonetheless, our findings illustrate the particular importance of the *F3*/*Tfpi* axis in thrombosis regulation (especially in the setting of *F5^L^*), as well as the identification of *Actr2* and the Arp2/3 complex as another potentially sensitive regulatory pathway for maintaining hemostatic balance.

## Materials and methods

Detailed description of mouse strains and procedures for ENU mutagenesis, breeding, genetic mapping and genotyping, Sanger and whole exome sequencing, estimation of *F3* allelic expression, generation of *Actr2* CRISPR/Cas9 targeted mice and cells, the *SURVEYOR nuclease assay*, and statistical data analyses are provided Supporting Information.

## Acknowledgements

This research was supported by NIH grants P01-HL057346 (to D. Ginsburg), R15-HL133907 and R01-HL135035 (to R. Westrick). R.J. Westrick was supported by the Oakland University Research Excellence Fund, an Aniara Diagnostica Coagulation Research Grant and American Heart Association Predoctoral Fellowship, Innovative Research, and Scientist Development grants. K. Tomberg was an International Fulbright Science and Technology Fellow and the recipient of an American Heart Association Predoctoral Fellowship Grant. M.A. Brake and A.J. Johnston were recipients of American Heart Association Undergraduate Fellowships. D. Ginsburg is a member of the University of Michigan Cancer Center. Research reported in this publication was supported by the National Cancer Institute of the National Institutes of Health under Award Number P30CA046592 by the use of the following Cancer Center Shared Resource(s): Transgenic Animal Models. The content is solely the responsibility of the authors and does not necessarily represent the official views of the National Institutes of Health. We gratefully acknowledge expertise of the Transgenic Animal Model Core staff of the University of Michigan’s Biomedical Research Core Facilities for assistance with this study. D. Ginsburg in an Investigator of the Howard Hughes Medical Institute.

## References

1. Silverstein MD, et al. (1998) Trends in the incidence of deep vein thrombosis and pulmonary embolism: a 25-year population-based study. Arch Intern Med 158(6):585–593.

2. Souto JC, et al. (2000) Genetic determinants of hemostasis phenotypes in Spanish families. Circulation 101(13):1546–1551.

3. Tregouet DA, et al. (2016) Is there still room for additional common susceptibility alleles for venous thromboembolism? J Thromb Haemost 14(9):1798–1802.

4. Dentali F, et al. (2012) Non-O blood type is the commonest genetic risk factor for VTE: results from a meta-analysis of the literature. Semin Thromb Hemost 38(5):535–548.

5. Rosendaal FR & Reitsma PH (2009) Genetics of venous thrombosis. J Thromb Haemost 7 Suppl 1:301–304.

6. Germain M, et al. (2015) Meta-analysis of 65,734 individuals identifies TSPAN15 and SLC44A2 as two susceptibility loci for venous thromboembolism. Am J Hum Genet 96(4):532–542.

7. Dahlback B (2008) Advances in understanding pathogenic mechanisms of thrombophilic disorders. Blood 112(1):19–27.

8. Lijfering WM, Rosendaal FR, & Cannegieter SC (2010) Risk factors for venous thrombosis – current understanding from an epidemiological point of view. Br J Haematol 149(6):824–833.

9. Clark JS, Adler G, Salkic NN, & Ciechanowicz A (2013) Allele frequency distribution of 1691G >A F5 (which confers Factor V Leiden) across Europe, including Slavic populations. J Appl Genet 54(4):441–446.

10. Westrick RJ & Ginsburg D (2009) Modifier genes for disorders of thrombosis and hemostasis. J Thromb Haemost 7 Suppl 1:132–135.

11. Evaluation of Genomic Applications in P & Prevention Working G (2011) Recommendations from the EGAPP Working Group: routine testing for Factor V Leiden (R506Q) and prothrombin (20210G>A) mutations in adults with a history of idiopathic venous thromboembolism and their adult family members. Genet Med 13(1):67–76.

12. De Stefano V, et al. (1999) The risk of recurrent deep venous thrombosis among heterozygous carriers of both factor V Leiden and the G20210A prothrombin mutation. N Engl J Med 341(11):801–806.

13. van Boven HH, et al. (1996) Factor V Leiden (FV R506Q) in families with inherited antithrombin deficiency. Thromb Haemost 75(3):417–421.

14. Cui J, et al. (2000) Spontaneous thrombosis in mice carrying the factor V Leiden mutation. Blood 96(13):4222–4226.

15. Eitzman DT, et al. (2002) Lethal perinatal thrombosis in mice resulting from the interaction of tissue factor pathway inhibitor deficiency and factor V Leiden. Circulation 105(18):2139–2142.

16. Cordes SP (2005) N-ethyl-N-nitrosourea mutagenesis: boarding the mouse mutant express. Microbiol Mol Biol Rev 69(3):426–439.

17. Moresco EM, Li X, & Beutler B (2013) Going forward with genetics: recent technological advances and forward genetics in mice. Am J Pathol 182(5):1462–1473.

18. Bull KR, et al. (2013) Unlocking the bottleneck in forward genetics using whole-genome sequencing and identity by descent to isolate causative mutations. PLoS Genet 9(1):e1003219.

19. Buchovecky CM, et al. (2013) A suppressor screen in Mecp2 mutant mice implicates cholesterol metabolism in Rett syndrome. Nat Genet 45(9):1013–1020.

20. Tchekneva EE, et al. (2007) A sensitized screen of N-ethyl-N-nitrosourea-mutagenized mice identifies dominant mutants predisposed to diabetic nephropathy. J Am Soc Nephrol 18(1):103–112.

21. Matera I, et al. (2008) A sensitized mutagenesis screen identifies Gli3 as a modifier of Sox10 neurocristopathy. Hum Mol Genet 17(14):2118–2131.

22. Carpinelli MR, et al. (2004) Suppressor screen in Mpl-/- mice: c-Myb mutation causes supraphysiological production of platelets in the absence of thrombopoietin signaling. Proc Natl Acad Sci U S A 101(17):6553–6558.

23. Bank I, et al. (2005) Elevated levels of FVIII:C within families are associated with an increased risk for venous and arterial thrombosis. J Thromb Haemost 3(1):79–84.

24. Nolan PM, et al. (2000) A systematic, genome-wide, phenotype-driven mutagenesis programme for gene function studies in the mouse. Nature Genetics 25(4):440–443.

25. Justice MJ, et al. (2000) Effects of ENU dosage on mouse strains. Mamm Genome 11(7):484–488.

26. Davis AP & Justice MJ (1998) An Oak Ridge legacy: the specific locus test and its role in mouse mutagenesis. Genetics 148(1):7–12.

27. Lusis AJ (2012) Genetics of atherosclerosis. Trends Genet 28(6):267–275.

28. Yoo YJ & Mendell NR (2008) The power and robustness of maximum LOD score statistics. Ann Hum Genet 72(Pt 4):566–574.

29. Kyrle PA, et al. (2004) The risk of recurrent venous thromboembolism in men and women. N Engl J Med 350(25):2558–2563.

30. Roach RE, et al. (2015) Sex difference in the risk of recurrent venous thrombosis: a detailed analysis in four European cohorts. J Thromb Haemost 13(10):1815–1822.

31. Vandenbroucke JP, et al. (2001) Oral contraceptives and the risk of venous thrombosis. N Engl J Med 344(20):1527–1535.

32. Justice MJ, Noveroske JK, Weber JS, Zheng B, & Bradley A (1999) Mouse ENU mutagenesis. Hum Mol Genet 8(10):1955–1963.

33. Rich JT, et al. (2010) A practical guide to understanding Kaplan-Meier curves. Otolaryngol Head Neck Surg 143(3):331–336.

34. Rotty JD, Wu C, & Bear JE (2013) New insights into the regulation and cellular functions of the ARP2/3 complex. Nat Rev Mol Cell Biol 14(1):7–12.

35. Rottner K, Hanisch J, & Campellone KG (2010) WASH, WHAMM and JMY: regulation of Arp2/3 complex and beyond. Trends in cell biology 20(11):650–661.

36. Veltman DM & Insall RH (2010) WASP family proteins: their evolution and its physiological implications. Molecular biology of the cell 21(16):2880–2893.

37. Falet H, et al. (2002) Importance of free actin filament barbed ends for Arp2/3 complex function in platelets and fibroblasts. Proc Natl Acad Sci U S A 99(26):16782–16787.

38. Vauti F, et al. (2007) Arp3 is required during preimplantation development of the mouse embryo. FEBS Lett 581(29):5691–5697.

39. Li Z, Kim ES, & Bearer EL (2002) Arp2/3 complex is required for actin polymerization during platelet shape change. Blood 99(12):4466–4474.

40. Koseoglu S, et al. (2015) VAMP-7 links granule exocytosis to actin reorganization during platelet activation. Blood 126(5):651–660.

41. Falet H, et al. (2002) Importance of free actin filament barbed ends for Arp2/3 complex function in platelets and fibroblasts. Proc Natl Acad Sci U S A 99(26):16782–16787.

42. Lek M, et al. (2016) Analysis of protein-coding genetic variation in 60,706 humans. Nature 536(7616):285–291.

43. McKusick VA (2007) Mendelian Inheritance in Man and its online version, OMIM. American journal of human genetics 80(4):588–604.

44. Beutler B (2017) Mutagenetix (TM).

45. Toomey JR, Kratzer KE, Lasky NM, Stanton JJ, & Broze GJ, Jr. (1996) Targeted disruption of the murine tissue factor gene results in embryonic lethality. Blood 88(5):1583–1587.

46. Huang ZF, Higuchi D, Lasky N, & Broze GJ, Jr. (1997) Tissue factor pathway inhibitor gene disruption produces intrauterine lethality in mice. Blood 90(3):944–951.

47. Bi L, et al. (1995) Targeted disruption of the mouse factor VIII gene produces a model of haemophilia A. Nat Genet 10(1):119–121.

48. Lange K, et al. (2013) Mendel: the Swiss army knife of genetic analysis programs. Bioinformatics 29(12):1568–1570.

49. Lander E & Kruglyak L (1995) Genetic dissection of complex traits: guidelines for interpreting and reporting linkage results. Nat Genet 11(3):241–247.

50. Mohlke KL, et al. (1999) Mvwf, a dominant modifier of murine von Willebrand factor, results from altered lineage-specific expression of a glycosyltransferase. Cell 96(1):111–120.

51. Tomberg K, et al. (2016) Spontaneous 8bp Deletion in Nbeal2 Recapitulates the Gray Platelet Syndrome in Mice. PLoS One 11(3):e0150852.

52. Li H & Durbin R (2009) Fast and accurate short read alignment with Burrows-Wheeler transform. Bioinformatics 25(14):1754–1760.

53. Garcia-Alcalde F, et al. (2012) Qualimap: evaluating next-generation sequencing alignment data. Bioinformatics 28(20):2678–2679.

54. DePristo MA, et al. (2011) A framework for variation discovery and genotyping using next-generation DNA sequencing data. Nat Genet 43(5):491–498.

55. Wang K, Li M, & Hakonarson H (2010) ANNOVAR: functional annotation of genetic variants from high-throughput sequencing data. Nucleic Acids Res 38(16):e164.

56. Ran FA, et al. (2013) Genome engineering using the CRISPR-Cas9 system. Nature protocols 8(11):2281–2308.

57. Brinster RL, Chen HY, Trumbauer ME, Yagle MK, & Palmiter RD (1985) Factors affecting the efficiency of introducing foreign DNA into mice by microinjecting eggs. Proc Natl Acad Sci U S A 82(13):4438–4442.

58. Yang H, et al. (2013) One-step generation of mice carrying reporter and conditional alleles by CRISPR/Cas-mediated genome engineering. Cell 154(6):1370–1379.

59. Sakurai T, Watanabe S, Kamiyoshi A, Sato M, & Shindo T (2014) A single blastocyst assay optimized for detecting CRISPR/Cas9 system-induced indel mutations in mice. BMC Biotechnol 14:69.

60. Therneau TM, Grambsch, P.M. (2000) Modeling survival data: extending the Cox model (Springer-Verlag New York) 1 Ed pp XIV, 350.

61. Brinkman EK, Chen T, Amendola M, & van Steensel B (2014) Easy quantitative assessment of genome editing by sequence trace decomposition. Nucleic Acids Res 42(22):e168.

62. Singh P, Schimenti JC, & Bolcun-Filas E (2015) A mouse geneticist’s practical guide to CRISPR applications. Genetics 199(1):1–15.

